# Insulin-like growth factor 1 signaling in the placenta requires endothelial nitric oxide synthase to support trophoblast function and normal fetal growth

**DOI:** 10.1101/2020.08.21.261420

**Authors:** Rebecca L Wilson, Weston Troja, Emily K Sumser, Alec Maupin, Kristin Lampe, Helen N Jones

## Abstract

Currently, there is no effective treatment for placenta dysfunction *in utero*. In a ligated mouse model of fetal growth restriction (FGR), nanoparticle-mediated *human insulin-like 1 growth factor* (*hIGF1*) gene delivery (NP-Plac1-hIGF1) increased *hIGF1* expression and maintained fetal growth. However, whether it can restore fetal growth remains to be determined. Using the endothelial nitric oxide synthase knockout (eNOS^−/−^) mouse model, a genetic model of FGR, we found that despite inducing expression of *hIGF1* in the placentas treated with NP-Plac1-hIGF1 (P=0.0425), FGR did not resolve. This was associated with no change to the number of fetal capillaries in the placenta labyrinth; an outcome increased with NP-Plac1-hIGF1 treatment in the ligated mouse model, despite increased expression of angiopoietin 1 (P=0.05) and suggested *IGF1* signaling in the placenta requires eNOS to modulate placenta angiogenesis. To further assess this hypothesis, we used the BeWo Choriocarcinoma cell line and human placenta explant cultures that were treated with NP-Plac1-hIGF1, oxidative stress was induced with hydrogen peroxide (H_2_O_2_) and NOS activity inhibited using the inhibitor L-NMMA. In both BeWo and explants, the protective effect of NP-Plac1-hIGF1 treatment against H_2_O_2_ induced cell death/lactate dehydrogenase release was prevented by eNOS inhibition (P=0.003 and P<0.0001, respectively). This was associated with an increase in mRNA expression of oxidative stress markers *HIF1α* (P<0.0001) and *ADAM10* (P=0.0002) in the NP-Plac1-hIGF1+H_2_O_2_+L-NMMA treated BeWo cells. These findings show for the first time the requirement of eNOS/NOS in IGF1 signaling in placenta cells that may have implications for placental angiogenesis and fetal growth.

## Introduction

Appropriate placentation underpins the existence of life. Numerous obstetric diseases including, miscarriage, stillbirth and fetal growth restriction (FGR) are associated with placenta dysfunction (22) as the placenta is the main interface between mother and fetus. In many cases, improper placental establishment results in inadequate utero-placental blood flow or impaired vascularization that ultimately reduces nutrient delivery and inevitably, fetal growth and development is compromised (4). While our understanding of the underlying causes and risk factors for placenta dysfunction is increasing (21), there is currently no effective *in utero* treatment for pregnant women who develop a placenta-related pregnancy complication. However, we are developing a polymer-based biodegradable nanoparticle that facilitates non-viral transgene delivery targeted to the placenta leading to enhanced placenta function (1, 31).

Insulin-like growth factor 1 (IGF1) is a single-chained polypeptide that through binding with the IGF1 receptor (IGF1R), is capable of affecting multiple cell functions including proliferation, differentiation, metabolism and apoptosis (25). Such actions are predominantly through the phosphoinositide-3 kinase (PI3K), protein kinase B (AKT) and mitogen-activated protein kinase (MAPK) signaling pathways. In terms of pregnancy and the placenta, IGF1 signaling is complex and involved in numerous processes vital to placenta function and fetal growth. Igf1 knockout (Igf1^−/−^) mice demonstrate fetal growth restriction (7, 17) and circulating levels of IGF1 are lower in human pregnancies complicated by FGR (2, 18). We and others previously demonstrated that increased placenta expression of *IGF1* in a surgically-induced mouse model of FGR, maintains fetal growth (1) through upregulation of glucose and amino acid transport mechanisms (13, 14) while adenoviral-mediated increase *hIGF1* in rabbits increases the weight of the runt fetus (15). Furthermore, systemic administration of IGF1 in the pregnant guinea pig from mid pregnancy increases fetal weight, placental uptake and transfer of nutrients and, fetal circulating levels of amino acids (23, 24). In terms of human placental function, increased *IGF1* expression in BeWo choriocarcinoma cells protects against increased cell death under oxidative stress conditions (31) and altogether, indicates that targeting IGF1 in the placenta may be effective in improving fetal growth.

Currently, there is no definitive way to predict, and therefore prevent, FGR and other major pregnancy complications. As such, potential therapies must be able to reverse the effects of existing placenta dysfunction. The endothelial nitric oxide synthase (eNOS^−/−^) knockout mouse model is a model of FGR, characterized with systemic vascular dysfunction, fetuses that are approximately 10% lighter in late pregnancy, and no gross placental malformations (9). eNOS is an enzyme which catalyzes the conversion of arginine to nitric oxide inducing vasodilation and as such, knocking out expression results in impaired uterine artery function and reduced amino acid transporter activity within the placenta (16). There is also increased generation of reactive oxygen species (ROS) within the eNOS^−/−^ placenta compared to the wildtype (16), an outcome that is often associated with placenta dysfunction in human pregnancies (5). As such, the eNOS^−/−^ mouse provides a suitable model to determine whether nanoparticle mediated h*IGF1* expression in the placenta could restore appropriate fetal growth.

## Materials and Methods

### Nanoparticle formation

Nanoparticles were formed by complexing a HPMA-DMEAMA copolymer with a plasmid encoding the human IGF1 gene under control of the PLAC1 promotor as previously described (1, 31).

### Animals

All animal procedures were approved by the Institutional Animal Care and Use Committee of Cincinnati Children’s Hospital Medical Center (Protocol number 2015-0078). Female eNOS^−/−^ on the C57Bl6 background (*n*=4) were obtained from Charles River Inc. and mated with male eNOS^−/−^; presence of a vaginal copulatory plug was designated GD0.5. At GD15.5, females underwent a laparotomy and all placenta from one uterine horn (*n*=2 left horn and *n*=2 right horn) were administered an intra-placenta (labyrinth region) injection of 20 μL NP-Plac1-hIGF1 (35 μg plasmid/placenta); all placentas situated in the opposite horn were not injected and designated as internal control. Animals were euthanized via CO_2_ asphyxiation 72 h after injection, cesarean-sections performed, and fetuses and placentas weighed. Each placenta was halved and snap-frozen and fixed in 4% w/v paraformaldehyde (PFA) and paraffin embedded for further analysis. Genomic DNA was extracted from each fetus and PCR used to determine fetal sex as previously described (30).

### Quantitative PCR (qPCR) in mouse placenta

For gene expression analysis, mouse placenta tissue was homogenized using the Qiagen TissueLyser II (*Qiagen*) with 5 mm beads in RLT buffer (*Qiagen*) containing 1% β-mercaptoethanol. RNA was extracted using the Qiagen RNAMini kit, including DNAse treatment, following the manufacturer’s instructions. 1 μg of RNA was the converted to cDNA using the High-capacity cDNA Reverse Transcription kit (*Applied Biosystems*). cDNA was diluted 1:100 and added to a reaction mix containing PowerUp SYBR Green (*Applied Biosystems*) as per manufacturer’s instructions and primers as outlined in supplemental table 1 (Table S1). qPCR was performed using the StepOne-Plus Real-Time PCR System (*Applied Biosystems*), and relative mRNA expression was calculated using the comparative CT method (20) with the StepOne Software v2.3 (*Applied Biosystems*) normalizing genes to *β-actin* and *RSP90*, which were confirmed to remain stable between internal control and NP-Plac1-hIGF1 treated samples.

### Histology and Immunohistochemistry in mouse placenta

For morphometric analysis, 4 placentas (2 internal control and 2 NP-Plac1-hIGF1 treated; 1 male and 1 female of each) from each dam were randomly chosen. 5 μm thick, full-faced sections were de-waxed and rehydrated according to standard protocols. Masson’s trichrome staining was used to visualize the labyrinth zone following standard protocol (*Sigma-Aldrich*) and labyrinth depth was calculated by averaging 3 random measurements (spanning the width of the section) from the base of the placenta to the labyrinth-junctional zone intersect. For immunohistochemical staining, 10 placentas (5 internal control and 5 NP-Plac1-hIGF1 treated) were randomly chosen for staining and antibodies and dilutions are provided in supplemental table 2 (Table S2). Antigen retrieval was performed by heating sections in 1x Targeting Retrieval Solution (*Dako*) for 30 mins at 95°C and 3% H_2_O_2_ was used to suppress endogenous peroxidase activity. Primary antibodies were diluted in 10% goat serum with 1% bovine serum albumin (BSA) and applied to slides overnight at 4°C. Negative controls omitting the primary antibody were included (Supplemental Fig 1). Secondary antibodies were diluted in 10% serum, 1% BSA and slides incubated for 1 h at room temperature. Antibody staining was then amplified using the Vector ABC kit (*Vector*) and detected using DAB (*Vector*) for a brown color or ImmPACT® VIP Substrate (*Vector*) for a purple color. Sections were counterstained with hematoxylin and imaged using the Nikon Eclipse 80i microscope and Nikon Elements Advanced Research software. Semi-quantitative analysis of IHC staining was performed using ImageJ software. First, color deconvolution was used to separate positive DAB/ImmPact® VIP substrate staining from hematoxylin and background noise. Mean grey intensity was then calculated on the DAB/ImmPact® VIP substrate color channel using the threshold function. This analysis was performed on 5 randomly chosen fields per placenta section and averaged to provide a result per placenta.

### BeWo and term placenta explant culture

Human BeWo choriocarcinoma cells (CCL-98; *American Type Culture Collection*) were cultured and treated in supplemented Ham’s F12 nutrient mix as previously described (31). Between cell passages 4 to 9 (*n*=6), 1 × 10^5^ BeWo cells/well were seeded onto 6-well plates, incubated for 4 h to allow attachment, and then treated with NP-Plac1-hIGF1 (8 μg plasmid/well) for 24 h to increase *hIGF1* mRNA expression. Oxidative stress was then induced by treatment with 400 μM H_2_O_2_ for 20 h (31) and NOS activity blocked by treatment with 1 mM L-NMMA (*Adipogen*) for 4 h. Cells were then collected and live/dead cell counts performed using trypan blue and a haemocytometer.

For human placenta explant experiments, de-identified, normal, term placentas (*n*=10, 4 male and 6 female) were approved and collected from University of Cincinnati Medical Center and The Christ Hospital. Villous explant samples (approximately 5 mm^3^) were dissected, washed and cultured, 6 explants/well of a 6-well plate, in supplemented 1:1 DMEM-Ham’s F12 media and cultured for 5 days prior to treatment as previously described (31). At 120 h, explants were treated for 24 h with NP-Plac1-hIGF1 (33 μg/well) followed by treatment with 400 μM H_2_O_2_ for 20 h to induce oxidative stress. NOS activity was then blocked by treatment with 1 mM L-NMMA for 4 h. Explant viability was assessed by measuring lactate dehydrogenase (LDH) release using the Pierce LDH Cytotoxicity Assay Kit (Thermo Scientific) follow manufacturer’s instructions and tissue was either snap-frozen for RNA and protein extractions or fixed in 4% w/v PFA and paraffin embedded for downstream analyses.

### BeWo and placental explant gene expression

For gene expression, BeWo cells were lysed using RLT buffer (*Qiagen*) and RNA extracted using the RNeasy mini kit (*Qiagen*) following manufacturer’s protocol. For the placenta explants, tissue was homogenized in Trizol (*Invitrogen*) with the aid of the Qiagen TissueLyser II and RNA purified following manufacturer’s instructions. Up to 10 μg extracted RNA was DNase treated using the TURBO DNA-free kit (*Applied Biosystems*) as per manufacturer’s protocol, and cDNA conversion and qPCR were performed as described above with primer sequences provided in supplemental table 1 (Table S1) and gene expression normalized to *β-actin* and *TBP*, which were confirmed to remain stable across treatments.

### Placental explant IHC

Immunohistochemical analysis of proteins was performed on explants from 5 placentas. 5 μm thick sections were treated as described above with antibodies and dilutions provided in supplemental table 2 (Table S2).

### Statistics

All statistical analyses were performed in R (v3.6.1)(26). For the eNOS^−/−^ mouse study, mixed effect linear modeling was used to calculate P-values with NP-Plac1-hIGF1 treatment selected as a fixed effect whilst maternal ID was selected as a random effect. For the *in vitro* BeWo and placenta fragment experiments, P-values were similarly calculated using mixed effect linear modeling; treatment was selected as a fixed effect whilst passage/placenta was selected as a random effect. Tukey’s post-hoc comparison of the least-square means was then used to compare between each treatment group. Data are represented as mean (SEM) or median (interquartile range: IQR) for mRNA and IHC data. Statistical significance was determined at P<0.05.

## Results

### Nanoparticle-mediated increased hIGF1 expression does not increase fetal growth in eNOS^−/−^ mice

Using qPCR analysis, there was no mRNA expression of *hIGF1* in the internal control placentas however, direct placenta injection of NP-Plac1-hIGF1 resulted in expression of *hIGF1* (median (IQR): 12 (1.3-29), P=0.0425). The increased expression of *hIGF1* did not change fetal or placental weight (Fig 1A and 1B; P=0.486 and P=0.893) nor placenta efficiency (Fig 1C; P=0.894). Fetal sex did not influence fetal or placental weight (Supplemental Fig 2).

**Figure 1.**
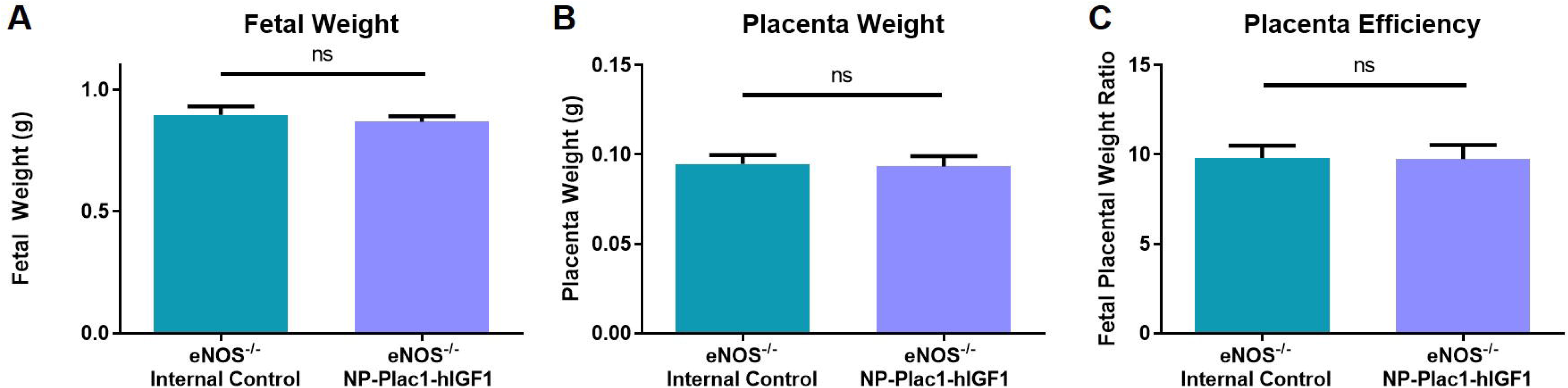
Effect of nanoparticle treatment (NP-Plac1-hIGF1) on fetal and placenta weights in the endothelial nitric oxide synthase knockout (eNOS^−/−^) mouse model. No difference in fetal (**A**) or placental (**B**) weight was observed with nanoparticle treatment when compared to non-injected internal control. There was also no difference in placenta efficiency (**C**). *n* = 4 dams (12 internal control and 11 NP-Plac1-hIGF1 fetuses/placentas). Data are mean±SEM.

### Nanoparticle-mediated hIGF1 expression in the eNOS^−/−^ mouse model did not increase labyrinth zone depth or the number of fetal capillaries

Surgical-ligation of the uterine artery branch resulted in reduced labyrinth zone (region of the placenta responsible for nutrient and waste exchange) thickness which was rescued with NP-Plac1-hIGF1 treatment (1). In the eNOS^−/−^ placentas, treatment with NP-Plac1-hIGF1 did not increase labyrinth zone depth compared to internal control (Fig. 2A; P=0.379). Maintained labyrinth zone depth with NP-Plac1-hIGF1 treatment in the surgical-ligation mouse model was associated with an maintenance in the number of fetal capillaries when compared to non-treated ligated placentas (mean (SEM) number of vessels per field: Sham=50.73 (2.15) vs. Ligated=37.47 (0.85) vs. Ligated+hIGF1=61.93 (7.16); P=0.02). However, a similar change to the number of fetal capillaries was not different in the NP-Plac1-hIGF1 treated eNOS^−/−^ placentas compared to internal control (Fig 2B; P=0.883). This was despite an increase in protein expression of angiopoietin 1 (Ang1) in the eNOS^−/−^ placenta treated with NP-Plac1-hIGF1 when compared to internal control placentas (Fig 2C; P=0.050). Protein expression of Ang2 did not differ with nanoparticle treatment (Fig 2D; P=0.651).

**Figure 2.**
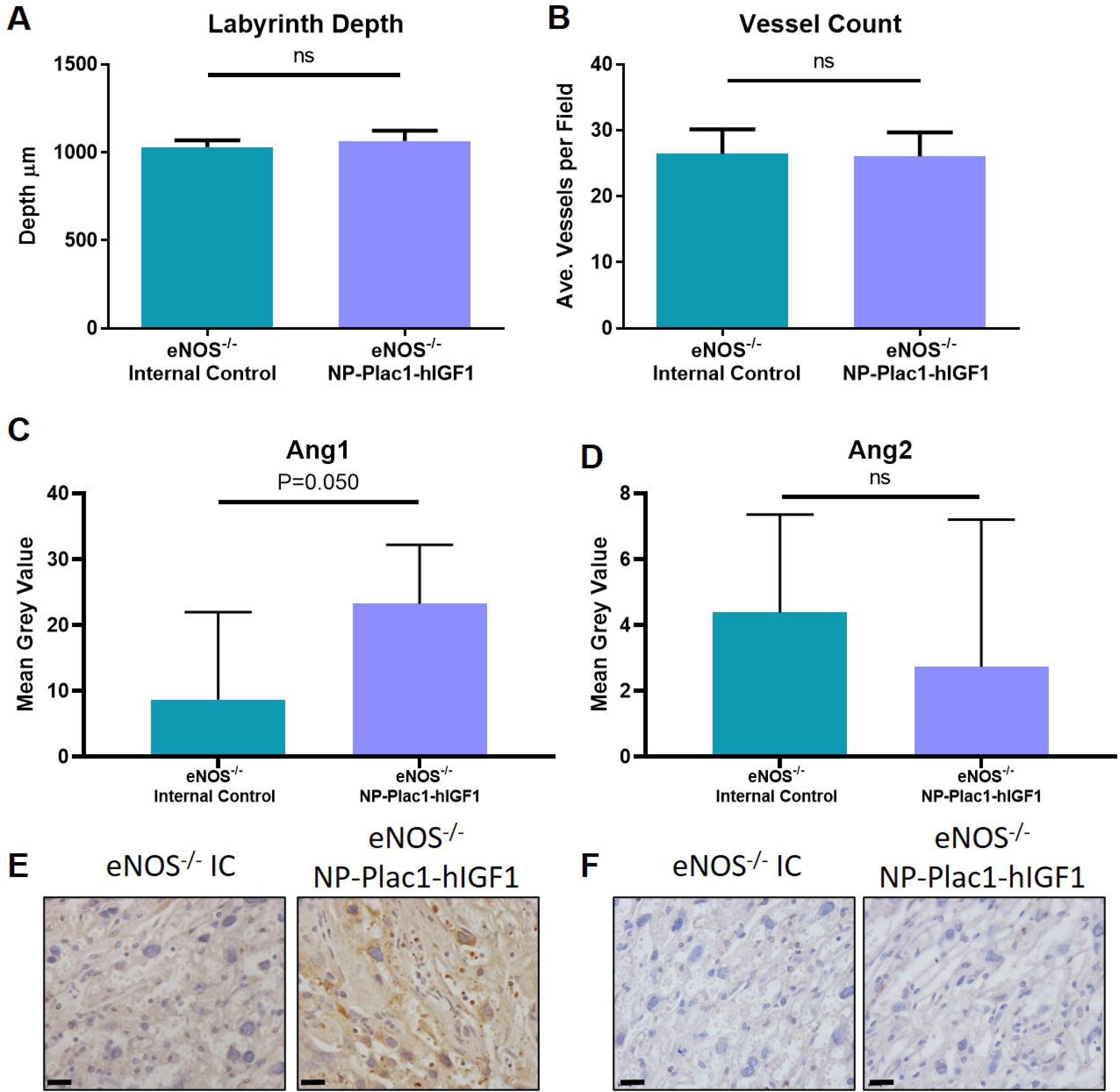
Effect of nanoparticle treatment (NP-Plac1-hIGF1) on placenta microstructure in the endothelial nitric oxide synthase knockout (eNOS^−/−^) mouse. NP-Plac1-hIGF1 treatment did not change the labyrinth zone depth (**A**) or the number of fetal vessels (**B**). Immunohistochemistry revealed increased expression of Angiopoietin 1 (Ang1) in the eNOS^−/−^ placentas treated with NP-Plac1-hIGF1 when compared to internal control (IC) placentas (**C**). However, protein expression of Ang2 was not different with nanoparticle treatment (**D**). Representative images of Ang1 (**E**) and Ang2 (**F**) stained with immunohistochemistry. Data for **A** and **B** are mean±SEM, n=4 pregnant eNOS^−/−^ mice (7-9 internal control and 8-9 NP-Plac1-hIGF1 placentas). Data for **C** and **D** are median±interquartile range, n=4 pregnant eNOS^−/−^ mice (5 internal control and 5 NP-Plac1-hIGF1 placentas). ns=non-significant. Scale bar=0.02 mm

### Nutrient transporter expression remains similar to internal controls despite increased hIGF1 expression in eNOS^−/−^ placentas

Maintenance of normal fetal growth in the surgically-induced mouse model of fetal growth restriction was facilitated by changes in placenta expression of glucose and amino acid transporters (14, 28). Compared to eNOS^−/−^ internal control placentas, NP-Plac1-hIGF1 treatment did not change the placenta mRNA expression of glucose transporter 1 and 8 (*Slc2a1* and *Slc2a8*, respectively) nor amino acid transporters *Slc38a1*, *Slc38a2*, *Slc7a5* and *Slc7a8* (Supplemental Table S3). Immunohistochemical analysis of Slc2a8 did however show increased protein expression with NP-Plac1-hIGF1 treatment compared to internal control (Fig 3A; P=0.038). There was however, no difference in protein expression of Slc2A1, Slc7a5 nor Slc38a2 between internal control and NP-Plac1-hIGF1 treated eNOS^−/−^ placentas (Fig 3B-D).

**Figure 3.**
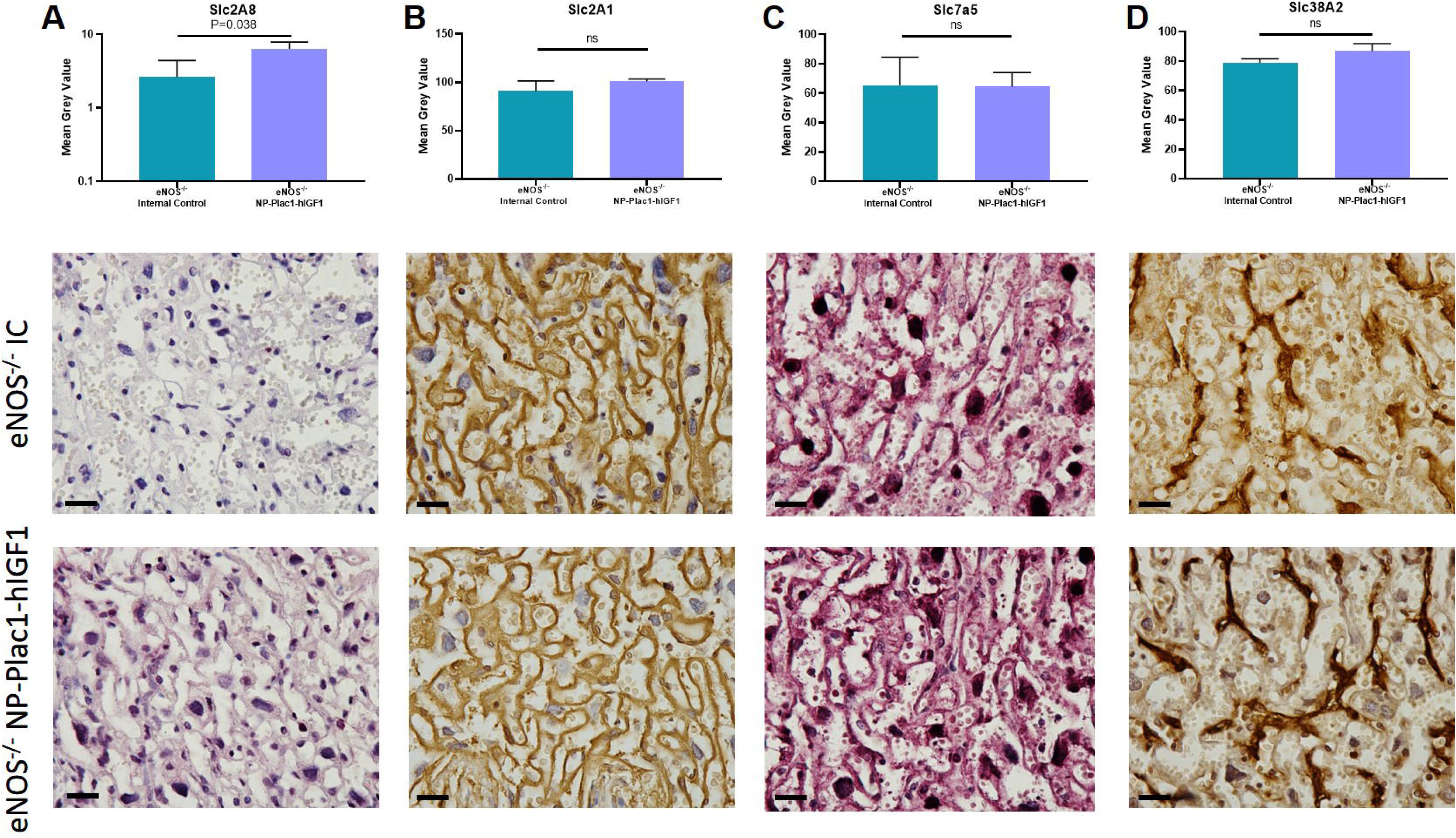
Protein expression of nutrient transporters in the endothelial nitric oxide synthase knockout (eNOS^−/−^) mouse placenta. Nanoparticle treatment (NP-Plac1-hIGF1) increased expression of Slc2A8 in the labyrinth of the eNOS^−/−^ placenta compared to non-injected internal control (IC) placentas as determined by immunohistochemistry (**A**). No difference in staining intensity of Slc2A1 (**B**), Slc7A5 (**C**), and Slc38A2 (**D**) was found. n=4 pregnant eNOS^−/−^ mice (5 internal control and 5 NP-Plac1-hIGF1 placentas). Data are median±interquartile range. Representative images for each transporter are presented below the graphs. Scale bar=0.02 mm

### IGF1 signaling requires NOS to protect against oxidative stress in BeWo cells

To support the findings in the eNOS^−/−^ mouse study, and further investigate our previous findings in BeWo cells in which NP-Plac1-hIGF1 treatment protected against increased cell death caused by H_2_O_2_ treatment (31), we treated BeWo cells with the NOS inhibitor L-NMMA. L-NMMA, did not affect nanoparticle-mediated increased *hIGF1* expression (Relative expression (median (IQR): untreated=1.4 (0.22-3.1) vs. NP-Plac1-hIGF1=2532 (742-4602) vs. L-NMMA=3.5 (0.86-6.2) vs. NP-Plac1-hIGF1+LNMMA=1371 (198-3969); P= 0.0087). Most importantly, the protective effect of NP-Plac1-hIGF1 treatment was not sustained when NOS activity was blocked as the percentage of dead cells was increased with H_2_O_2_+L-NMMA and NP-Plac1-hIGF1+H_2_O_2_+L-NMMA treatment. This was not due to L-NMMA treatment as there was no difference in the percentage of dead cells with L-NMMA treatment alone (mean (SEM) %: untreated=16 (3.2) vs. L-NMMA=17 (2.9) vs. NP-Plac1-hIGF1+LNMMA=18 (4.5) vs. H_2_O_2_+L-NMMA=32 (6.1) vs. NP-Plac1-hIGF1+H_2_O_2_+L-NMMA=28 (4); P=0.003). Gene expression analysis of markers of oxidative stress revealed increased mRNA expression of *hypoxia inducing factor 1α* (*HIF1α*) and *ADAM10* in the NP-Plac1-hIGF1+H_2_O_2_+L-NMMA cells compared to other treatments (Fig 4A; P<0.0001 and Fig 4B; P=0.0002, respectively). Interestingly, L-NMMA treatment increased the expression of *P53* in BeWo cells (Fig 4C; P<0.0001) whilst all cells treated with H_2_O_2_ had increased *HIF2α* expression (Fig 4C; P=0.038). Expression of *superoxide dismutase 1* (*SOD1*) and *SOD2* was not different with any treatment (Fig 4e & F).

**Figure 4.**
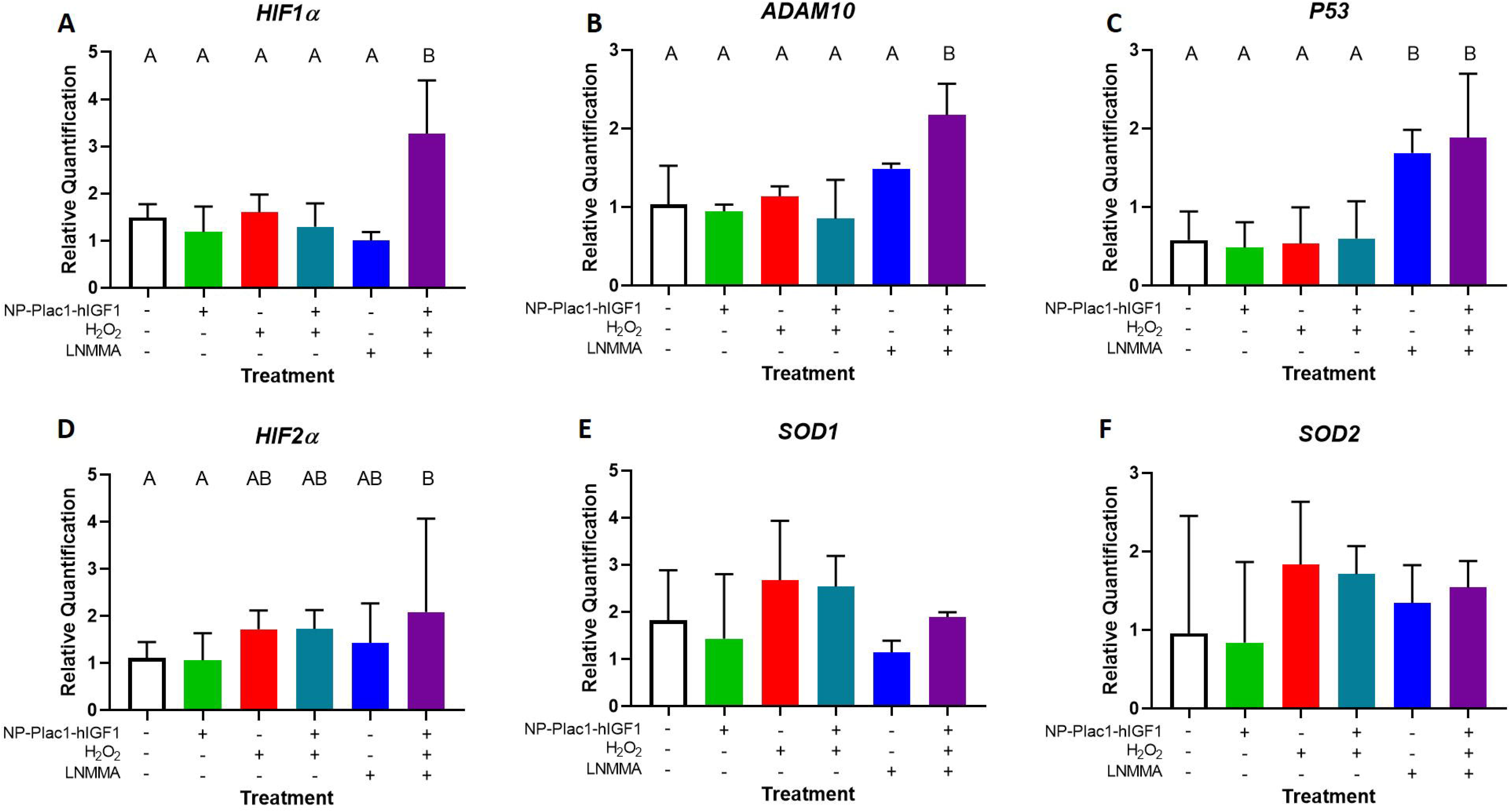
Effect of nanoparticle (NP-Plac1-hIGF1), hydrogen peroxide (H_2_O_2_), and nitric oxide synthase (NOS) inhibitor (L-NMMA) treatment in BeWo cells. NP-Plac1-hIGF1+H_2_O_2_+L-NMMA treatment increased mRNA expression of *HIF1α* (**A**) and *ADAM10* (**B**) compared to all other treatments. Treatment with L-NMMA had increased mRNA expression of *P53* compared to cells not treated with L-NMMA (**C**) whilst expression of *HIF2α* also appeared to change with L-NMMA and H_2_O_2_ treatment (**D**). mRNA expression of *SOD1* (**E**) and *SOD2* (**F**) was not different across the treatment groups. Data are median±interquartile range of 6 independent experimental replicates. Different letters denote significant differences of P<0.05.

### NOS inhibition effects nanoparticle mediated changes to human placenta explant function

To assess findings in the context of the placental milieu, we treated human placenta explants collected from term pregnancies with NP-Plac1-hIGF1, H_2_O_2_ and L-NMMA. All explants treated with nanoparticle had increased *hIGF1* mRNA expression (Fig 5; P=0.0026). Given that IGF1 is a secreted protein and not easily measured with standard protocols (Supplemental Fig 3), we assessed downstream changes to explant nutrient transporter expression which would only occur if increased *hIGF1* mRNA expression resulted in increased hIGF1 signaling. Consistent with prior *in vivo* and *in vitro* investigations (13–15, 31), treatment of explants with NP-Plac1-hIGF1 resulted in increased expression of SLC2A8 (Fig 6A; P=0.011), SLC7A5 (Fig 6B; P=0.001) and SLC38A2 (Fig 6C; P=0.0002). L-NMMA treatment did not affect the increase caused by NP-Plac1-hIGF1 treatment nor was there a difference in SLC2A1 protein expression (Fig 6D).

**Figure 5.**
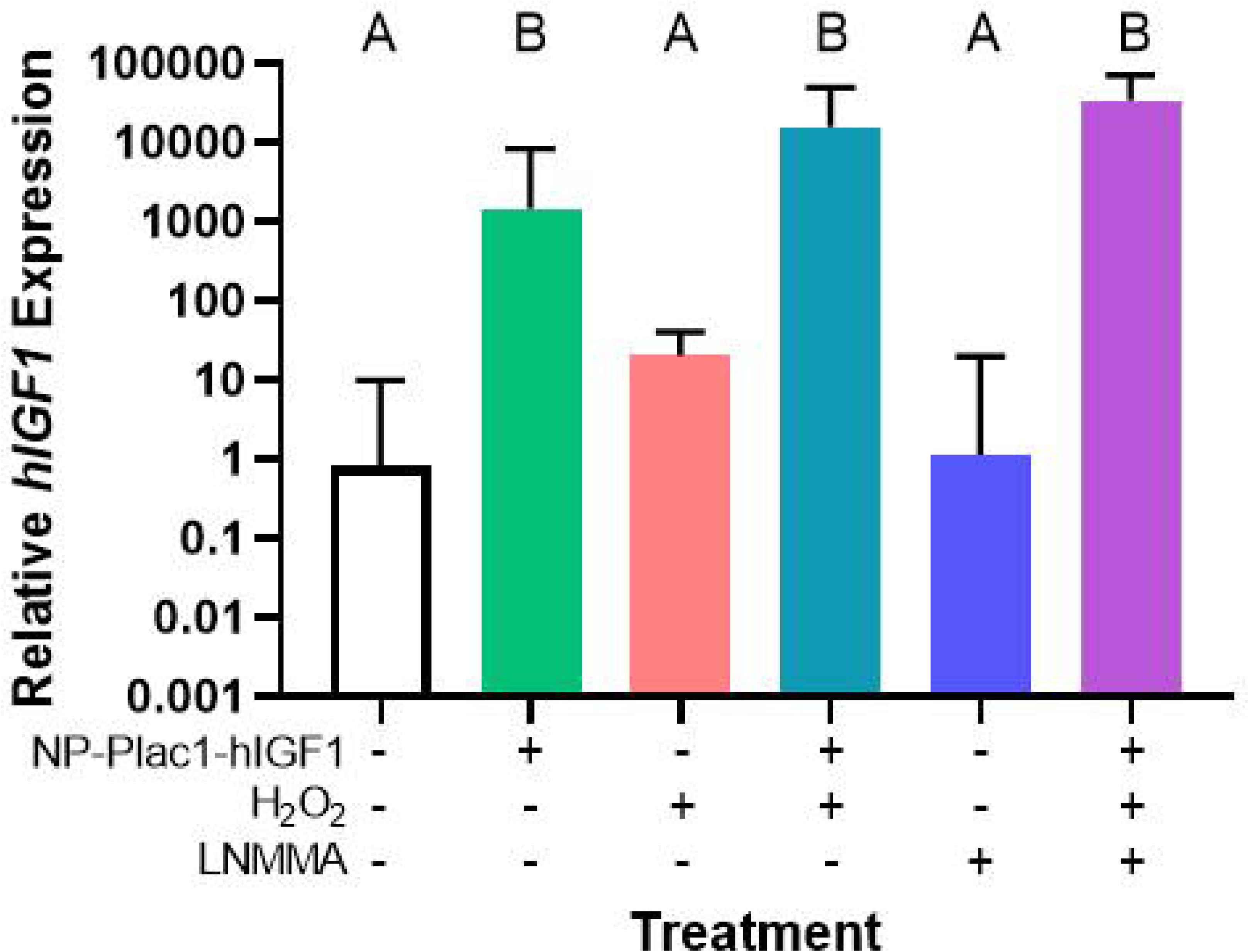
Effect of nanoparticle (NP-Plac1-hIGF1), hydrogen peroxide (H_2_O_2_) and, nitric oxide synthase (NOS) inhibitor (L-NMMA) treatment on human *Insulin-like growth factor 1* (*hIGF1*) mRNA expression in term placenta explants. All explants treated with NP had increased *Insulin-like growth factor* (*hIGF1*) mRNA expression. Data are median±interquartile range of n=5-10 term placentas. Different letters denote significant differences of P<0.05.

**Figure 6.**
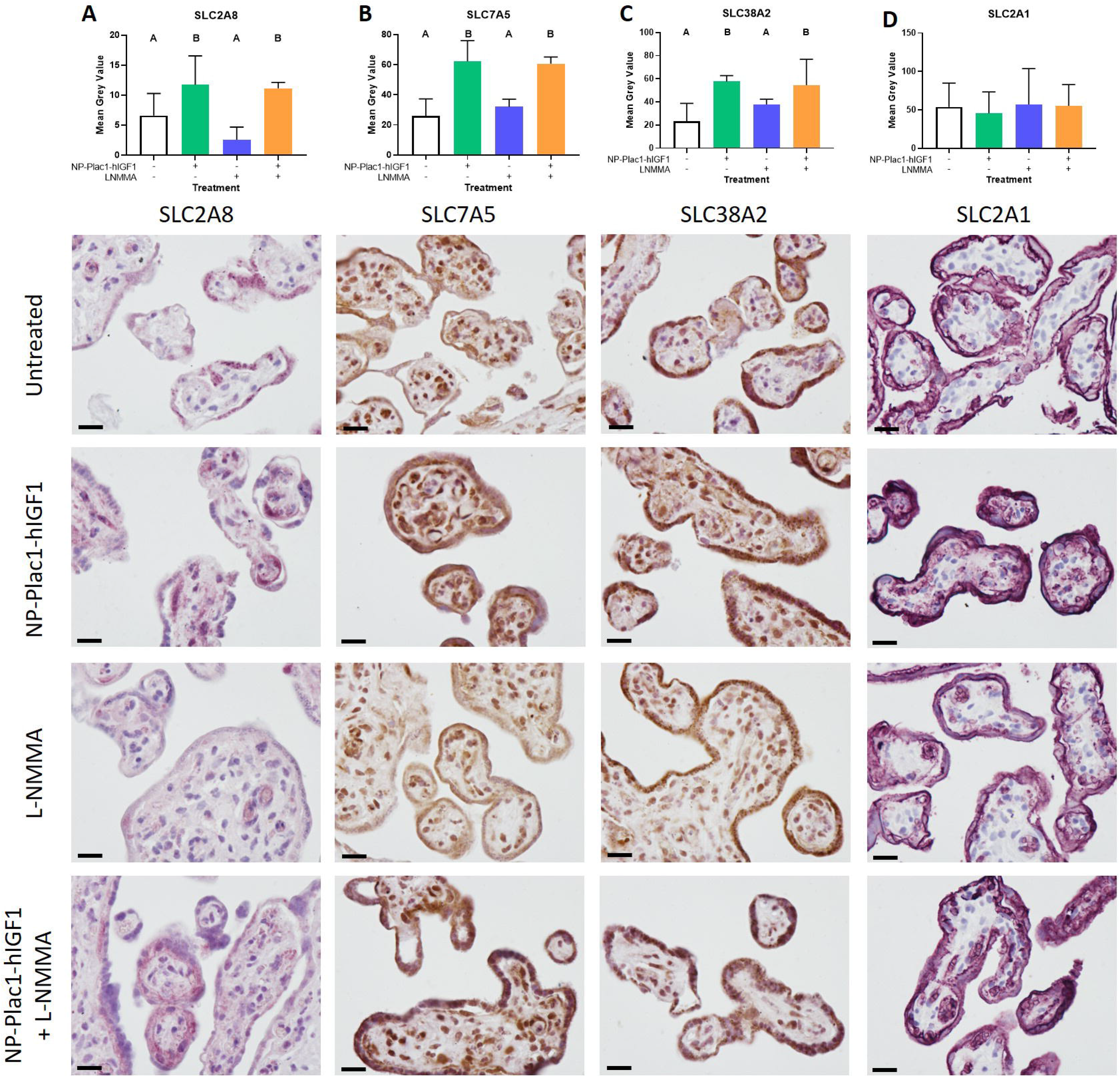
Protein expression of nutrient transporters in human placenta explants treated with nanoparticle (NP-Plac1-hIGF1) and nitric oxide synthase (NOS) inhibitor L-NMMA. Immunohistochemistry revealed that nanoparticle treatment increased expression of SLC2A8 (**A**), SLC7A5 (**B**) and SLC38A2 (**C**). However, there was no difference in protein expression of SLC2A1 (**D**) was found with nanoparticle treatment. There was also no clear effect of LNMMA treatment on any of the transporters. Data are median±interquartile range and representative images of explant cultures from 5 different term placentas for each transporter are presented below the graphs. Different letters denote significant difference of P<0.05. Scale bar=0.02 mm

LDH release was increased with H_2_O_2_ treatment but not when explants were also treated with NP-Plac1-hIGF1 which remained at a similar level to untreated and NP-Plac1-hIGF1 only treatment (Fig 7A; P=<0.0001). Most importantly, LDH release was not affected by L-NMMA treatment and increased in the explants treated with NP-Plac1-hIGF1, H_2_O_2_ and L-NMMA when compared to explants not treated with H_2_O_2_ (Fig 7A). In further understanding this, we analyzed explant mRNA expression of markers of oxidative stress but found only the *BAX:BCL2* ratio to be increased in explants treated with NP-Plac1-hIGF1+ H_2_O_2_+L-NMMA (Fig 7B; P=0.0062). There was no difference in the mRNA expression of *HIF1α*, *HIF2α*, *SOD1* nor *SOD2* (Fig 7C-F).

**Figure 7.**
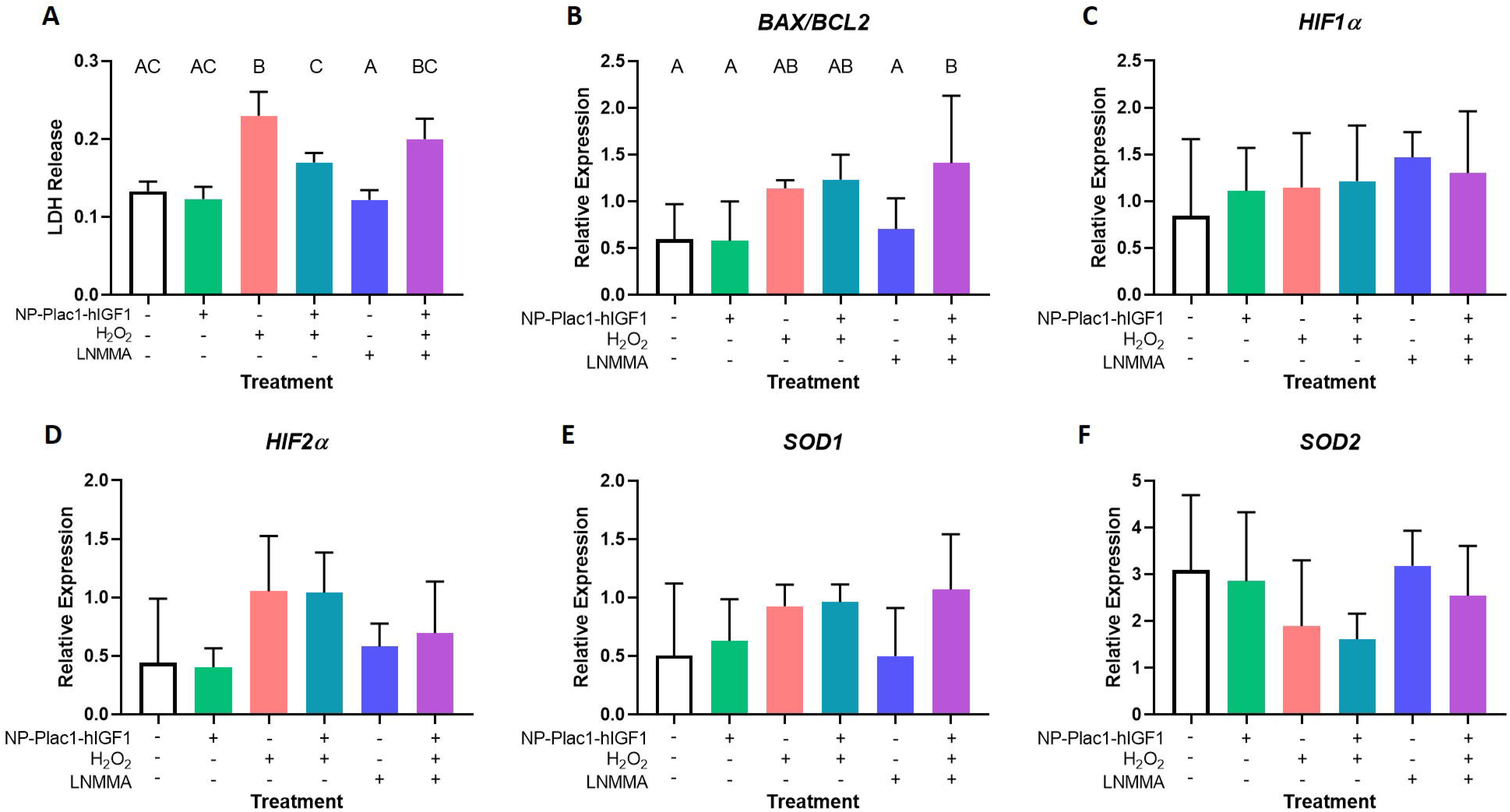
Effect of nanoparticle (NP-Plac1-hIGF1), hydrogen peroxide (H_2_O_2_) and, nitric oxide synthase (NOS) inhibitor (L-NMMA) treatment in term placenta explants. Treatment with H_2_O_2_ increased LDH release, was protected against with NP-Plac1-hIGF1 treatment except when NOS activity was blocked by L-NMMA (**A**). Analysis of mRNA expression showed an increase in the *BAX:BCL2* ratio in explants treated with NP-Plac1-hIGF1+H_2_O_2_+L-NMMA when compared to untreated, NP-Plac1-hIGF1 and L-NMMA only treated explants (**B**). There was no difference in mRNA expression of *Hypoxia inducing factor 1α* (*HIF1α*; **C**), *HIF2α* (**D**), *Superoxide dismutase 1* (*SOD1*; **E**) or *SOD2* (**F**). Data are mean+SEM (**A**) and median+interquartile range (**B**-**F**) of n=5-10 term placentas. Different letters denote significant differences of P<0.05.

## Discussion

This study shows for the first time that IGF1 signaling in placenta trophoblast cells relies on eNOS as an intermediate and has implications for placenta angiogenesis and fetal growth. The ability for increased *IGF1* expression in the placenta to maintain fetal growth is well established (1), however, the ability to restore fetal growth in a model of pre-existing FGR remains unknown. Using the transgenic eNOS^−/−^ mouse model, we found that despite nanoparticle-mediated *hIGF1* expression and increased expression of Ang1 and nutrient Slc2A8, there was no change to fetal growth. Hence identifying the novel requirement of eNOS for IGF1 signaling in trophoblast cells. This was corroborated in the *in vitro* experiments in which, nanoparticle-mediated increased *IGF1* protected against increased cellular stress but was blocked by treatment with the NOS inhibitor L-NMMA. As such, the current study provides crucial knowledge about IGF1 signaling in the placenta not previously demonstrated, as well as implications for further developing methods to optimize fetal growth in utero.

The initial aim of this study was to determine whether nanoparticle-mediated increased *hIGF1* expression could enhance placenta function and restore appropriate fetal growth in a transgenic model of pre-existing FGR. While the eNOS^−/−^ placenta was capable of nanoparticle uptake and increased transgene expression, this ultimately had no effect on either placental or fetal growth. Further analysis indicated this may be due to the inability for IGF1 to promote placenta angiogenesis. Despite increased Ang1 protein expression in the NP-Plac1-hIGF1 treated placentas, there was no change in fetal vessel number in the labyrinth of the placenta. In the ligated mouse model, in which fetal growth was maintained with nanoparticle treatment, increased *IGF1* expression increased the number of fetal vessels within the placenta. This outcome would be positive for fetal growth and is complementary to others that have shown that IGF1 increases the number of micro-vessels in cultured rat aortic explants and IGF1 receptor binding increases tube formation of endothelial progenitor cells (10, 19). Ang1 is an integral mediator of angiogenesis and controls numerous signaling pathways that support the formation and function of vessels (8). Furthermore, Ang1 has been shown to stimulate eNOS phosphorylation *in vitro* (6). Based on our results, it is likely that nanoparticle treatment results in increased expression of Ang1 but is unable to support increased placental angiogenesis due to a lack of eNOS and thus results in no change in fetal growth in the eNOS^−/−^ mouse. Overall, this outcome indicates a synergistic relationship between IGF1 and eNOS signaling pathways in placental trophoblast not previously shown but worth further investigating.

In non-trophoblast studies, IGF-1 has is known to increase eNOS phosphorylation and activity in endothelial progenitor cells and vascular smooth muscle cells (12, 27). Furthermore, IGF1 protects endothelial progenitor cells from increased apoptosis caused by oxidized low-density lipoproteins; an effect abolished by the presence of L-NAME, another NOS inhibitor (29). In our study, a similar outcome was observed as increased *hIGF1* expression protected BeWo cells against increased cell death caused by oxidative stress but was prevented when NOS activity was inhibited. Similarly, this affect was observed with LDH release in the placenta explant cultures. In understanding this further, we observed increased mRNA expression of oxidative stress markers *HIF1α* and *ADAM10*. Changes to *HIF1α* and *ADAM10* expression are rapid and occur shortly after the induction of cellular stress (3, 32). At the conclusion of the BeWo cell culture model, expression of *HIF1α* and *ADAM10* was increased in the cells treated with NP-Plac1-hIGF1+H_2_O_2_+L-NMMA but not in the H_2_O_2_ or NP-Plac1-hIGF1+H_2_O_2_ treated cells. This could be explained by experimental timing; H_2_O_2_ treatment was administered 24 h prior and any changes to *HIF1α* and *ADAM10* expression caused by the increased oxidative stress conditions may have already occurred. As such, it could be hypothesized that nanoparticle-mediated increased *hIGF1* expression suppresses increased *HIF1α* and *ADAM10* caused by H_2_O_2_ treatment but when NOS activity is inhibited this suppression is prevented allowing *HIF1α* and *ADAM10* to increase and result in cell death. Whilst further experiments, beyond the scope of this manuscript are required to confirm this, it does further strengthen the understanding that IGF1 signaling in trophoblast cells requires eNOS/NOS signaling.

Explants were used to further characterize the outcomes of nanoparticle treatment and NOS inhibition in a culture system better reflective of placental physiology. However, unlike in the BeWo cells, we did not find any significant changes in the mRNA expression of oxidative stress markers except for the *BAX:BCL2* ratio which was elevated with H_2_O_2_ treatment. This could be explained by a number of factors including the presence of multiple cell types and experimental timing. The placenta explant surface is composed of the multinucleated syncytiotrophoblast, a terminally differentiated layer replenished by fusion of the underlying cytotrophoblasts (11). Together these trophoblast subpopulations encapsulate the mesenchymal core which contains the fetal capillaries. We have previously shown that nanoparticle mediated transgene expression is confined to the syncytiotrophoblasts at 72 hours post treatment (31) and therefore most likely to be exhibiting the most significant changes with nanoparticle, H_2_O_2_ and L-NMMA treatment. The short duration of L-NMMA treatment may also influence outcomes of the study and likely the reason why no statistically significance difference in LDH release was observed between the NP-Plac1-hIGF1+H_2_O_2_ and NP-Plac1-hIGF1+H_2_O_2_+L-NMMA. Finally, it is possible outcomes in the explant study may be influenced by inhibition of all three NOS isoforms as L-NMMA inhibits the active site of eNOS, inducible NOS and neuronal NOS. Whilst further investigations beyond the scope of this study are required, particularly focusing on individual cell responses as well as the contribution of the different NOS isoforms, such data provides a solid framework for continuing to elucidate the mechanisms between IGF1 signaling, eNOS and placental angiogenesis.

## Conclusions

In summary, our study is the first to provide strong evidence that eNOS is an important intermediate in the IGF1 signaling pathway in the placenta. We propose that eNOS is required for IGF1 to modulate placental angiogenesis as well as protect trophoblast cells against cellular stress and provides justification as to why nanoparticle treatment in the eNOS^−/−^ mouse model was unable to restore fetal growth. These studies provide a framework for continuing to determine the mechanistic link between IGF1 and eNOS/NOS signaling in trophoblast and how this affects placental development and function and fetal growth.

## Non-standard abbreviations

FGR: fetal growth restriction
hIGF1: human insulin-like growth factor 1
NP: nanoparticle
NOS: nitric oxide synthase
eNOS: endothelial nitric oxide synthase
L-NMMA: N^5^-[imino(methylamino)methyl]-L-ornithine, monoacetate
ANG1: angiopoietin 1
ANG2: angiopoietin 2
HIF1α: hypoxia inducing factor 1α
HIF2α: hypoxia inducing factor 2α
SOD1: superoxide dismutase 1
SOD2: superoxide dismutase 2
PGF: placenta growth factor
PDGFβ: platelet-derived growth factor β
PDGFβR: platelet-derived growth factor β receptor
VEGFβ: vascular endothelial growth factor β

## Declarations

The authors declare no conflicts of interest

RLW designed and performed experiments, analyzed the data and wrote the manuscript. WT, EKS, AM and KL performed experiments, analyzed the data and edited the manuscript. HNJ conceived the study, and edited the manuscript. All authors approved the final manuscript.

This study was funded by Eunice Kennedy Shriver National Institute of Child Health and Human Development (NICHD) award R01HD090657 (HNJ).

## Notes

### Competing Interest Statement

The authors have declared no competing interest.

## References

1. Abd Ellah N, Taylor L, Troja W, Owens K, Ayres N, Pauletti G, and Jones H. Development of Non-Viral, Trophoblast-Specific Gene Delivery for Placental Therapy. PloS one 10: e0140879, 2015.

2. Ashton IK, Zapf J, Einschenk I, and MacKenzie IZ. Insulin-like growth factors (IGF) 1 and 2 in human foetal plasma and relationship to gestational age and foetal size during midpregnancy. Acta Endocrinol (Copenh) 110: 558–563, 1985.

3. Bergeron M, Yu AY, Solway KE, Semenza GL, and Sharp FR. Induction of hypoxia-inducible factor-1 (HIF-1) and its target genes following focal ischaemia in rat brain. Eur J Neurosci 11: 4159–4170, 1999.

4. Brosens I, Pijnenborg R, Vercruysse L, and Romero R. The “Great Obstetrical Syndromes” are associated with disorders of deep placentation. Am J Obstet Gynecol 204: 193–201, 2011.

5. Burton GJ, Yung HW, Cindrova-Davies T, and Charnock-Jones DS. Placental endoplasmic reticulum stress and oxidative stress in the pathophysiology of unexplained intrauterine growth restriction and early onset preeclampsia. Placenta 30 Suppl A: S43–48, 2009.

6. Chen JX, Lawrence ML, Cunningham G, Christman BW, and Meyrick B. HSP90 and Akt modulate Ang-1-induced angiogenesis via NO in coronary artery endothelium. J Appl Physiol (1985) 96: 612–620, 2004.

7. Constancia M, Hemberger M, Hughes J, Dean W, Ferguson-Smith A, Fundele R, Stewart F, Kelsey G, Fowden A, Sibley C, and Reik W. Placental-specific IGF-II is a major modulator of placental and fetal growth. Nature 417: 945–948, 2002.

8. Dunk C, Shams M, Nijjar S, Rhaman M, Qiu Y, Bussolati B, and Ahmed A. Angiopoietin-1 and angiopoietin-2 activate trophoblast Tie-2 to promote growth and migration during placental development. Am J Pathol 156: 2185–2199, 2000.

9. Hefler LA, Tempfer CB, Moreno RM, O’Brien WE, and Gregg AR. Endothelial-derived nitric oxide and angiotensinogen: blood pressure and metabolism during mouse pregnancy. Am J Physiol Regul Integr Comp Physiol 280: R174–182, 2001.

10. Humpert PM, Djuric Z, Zeuge U, Oikonomou D, Seregin Y, Laine K, Eckstein V, Nawroth PP, and Bierhaus A. Insulin stimulates the clonogenic potential of angiogenic endothelial progenitor cells by IGF-1 receptor-dependent signaling. Mol Med 14: 301–308, 2008.

11. Huppertz B, and Borges M. Placenta trophoblast fusion. Methods Mol Biol 475: 135–147, 2008.

12. Isenovic ER, Meng Y, Divald A, Milivojevic N, and Sowers JR. Role of phosphatidylinositol 3-kinase/Akt pathway in angiotensin II and insulin-like growth factor-1 modulation of nitric oxide synthase in vascular smooth muscle cells. Endocrine 19: 287–292, 2002.

13. Jones H, Crombleholme T, and Habli M. Regulation of amino acid transporters by adenoviral-mediated human insulin-like growth factor-1 in a mouse model of placental insufficiency in vivo and the human trophoblast line BeWo in vitro. Placenta 35: 132–138, 2014.

14. Jones HN, Crombleholme T, and Habli M. Adenoviral-mediated placental gene transfer of IGF-1 corrects placental insufficiency via enhanced placental glucose transport mechanisms. PLoS One 8: e74632, 2013.

15. Keswani SG, Balaji S, Katz AB, King A, Omar K, Habli M, Klanke C, and Crombleholme TM. Intraplacental gene therapy with Ad-IGF-1 corrects naturally occurring rabbit model of intrauterine growth restriction. Hum Gene Ther 26: 172–182, 2015.

16. Kusinski LC, Stanley JL, Dilworth MR, Hirt CJ, Andersson IJ, Renshall LJ, Baker BC, Baker PN, Sibley CP, Wareing M, and Glazier JD. eNOS knockout mouse as a model of fetal growth restriction with an impaired uterine artery function and placental transport phenotype. Am J Physiol Regul Integr Comp Physiol 303: R86–93, 2012.

17. Liu JP, Baker J, Perkins AS, Robertson EJ, and Efstratiadis A. Mice carrying null mutations of the genes encoding insulin-like growth factor I (Igf-1) and type 1 IGF receptor (Igf1r). Cell 75: 59–72, 1993.

18. McIntyre HD, Serek R, Crane DI, Veveris-Lowe T, Parry A, Johnson S, Leung KC, Ho KK, Bougoussa M, Hennen G, Igout A, Chan FY, Cowley D, Cotterill A, and Barnard R. Placental growth hormone (GH), GH-binding protein, and insulin-like growth factor axis in normal, growth-retarded, and diabetic pregnancies: correlations with fetal growth. J Clin Endocrinol Metab 85: 1143–1150, 2000.

19. Nicosia RF, Nicosia SV, and Smith M. Vascular endothelial growth factor, platelet-derived growth factor, and insulin-like growth factor-1 promote rat aortic angiogenesis in vitro. Am J Pathol 145: 1023–1029, 1994.

20. Pfaffl MW. A new mathematical model for relative quantification in real-time RT-PCR. Nucleic Acids Res 29: e45, 2001.

21. Redman CW, and Staff AC. Preeclampsia, biomarkers, syncytiotrophoblast stress, and placental capacity. Am J Obstet Gynecol 213: S9 e1, S9-11, 2015.

22. Roberts CT. IFPA Award in Placentology Lecture: Complicated interactions between genes and the environment in placentation, pregnancy outcome and long term health. Placenta 31 Suppl: S47–53, 2010.

23. Sferruzzi-Perri AN, Owens JA, Pringle KG, Robinson JS, and Roberts CT. Maternal insulin-like growth factors-I and - II act via different pathways to promote fetal growth. Endocrinology 147: 3344–3355, 2006.

24. Sferruzzi-Perri AN, Owens JA, Standen P, Taylor RL, Robinson JS, and Roberts CT. Early pregnancy maternal endocrine insulin-like growth factor I programs the placenta for increased functional capacity throughout gestation. Endocrinology 148: 4362–4370, 2007.

25. Sferruzzi-Perri AN, Sandovici I, Constancia M, and Fowden AL. Placental phenotype and the insulin-like growth factors: resource allocation to fetal growth. J Physiol 595: 5057–5093, 2017.

26. Team RC. R: A Language and Environment for Statistical Computing. Vienna, Austria: R Foundation for Statistical Computing, 2019.

27. Thum T, Fleissner F, Klink I, Tsikas D, Jakob M, Bauersachs J, and Stichtenoth DO. Growth hormone treatment improves markers of systemic nitric oxide bioavailability via insulin-like growth factor-I. J Clin Endocrinol Metab 92: 4172–4179, 2007.

28. Troja W, Kil K, Klanke C, and Jones HN. Interaction between human placental microvascular endothelial cells and a model of human trophoblasts: effects on growth cycle and angiogenic profile. Physiol Rep 2: e00244, 2014.

29. Wen HJ, Liu GF, Xiao LZ, and Wu YG. Involvement of endothelial nitric oxide synthase pathway in IGF1 protects endothelial progenitor cells against injury from oxidized LDLs. Mol Med Rep 19: 660–666, 2019.

30. Wilson RL, Buckberry S, Spronk F, Laurence JA, Leemaqz S, O’Leary S, Bianco-Miotto T, Du J, Anderson PH, and Roberts CT. Vitamin D Receptor Gene Ablation in the Conceptus Has Limited Effects on Placental Morphology, Function and Pregnancy Outcome. PLoS One 10: e0131287, 2015.

31. Wilson RL, Owens K, Sumser EK, Fry MV, Stephens KK, Chuecos M, Carrillo M, Schlabritz-Loutsevitch N, and Jones HN. Nanoparticle mediated increased insulin-like growth factor 1 expression enhances human placenta syncytium function. Placenta 93: 1–7, 2020.

32. Yang Y, Wang Y, Zeng X, Ma XJ, Zhao Y, Qiao J, Cao B, Li YX, Ji L, and Wang YL. Self-control of HGF regulation on human trophoblast cell invasion via enhancing c-Met receptor shedding by ADAM10 and ADAM17. J Clin Endocrinol Metab 97: E1390–1401, 2012.

