## Supplemental Material for "Insulin-like growth factor 1 signaling in the placenta requires endothelial nitric oxide synthase to support trophoblast function and normal fetal growth"

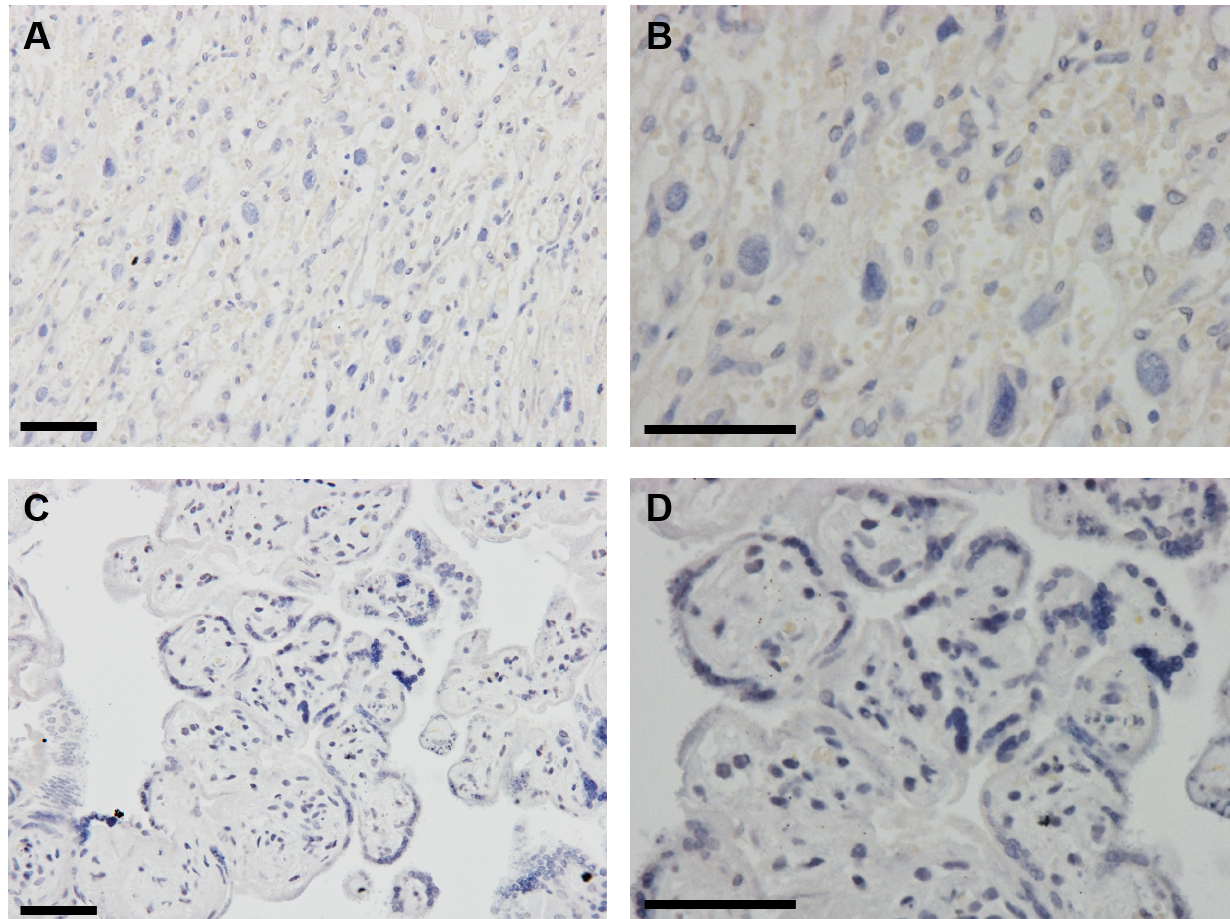

**Supplemental Figure 1** Representative images of immunohistochemistry (IHC) negative controls. For all antibodies stained using IHC, negative controls were included and showed no positive staining for both eNOS^-/-^ mouse placenta tissue (**A** & **B**) and human explant tissue (**C** & **D**). Scale bar = 0.05 mm. **A** and **C** captured at 20x, **B** and **D** captured at 40x

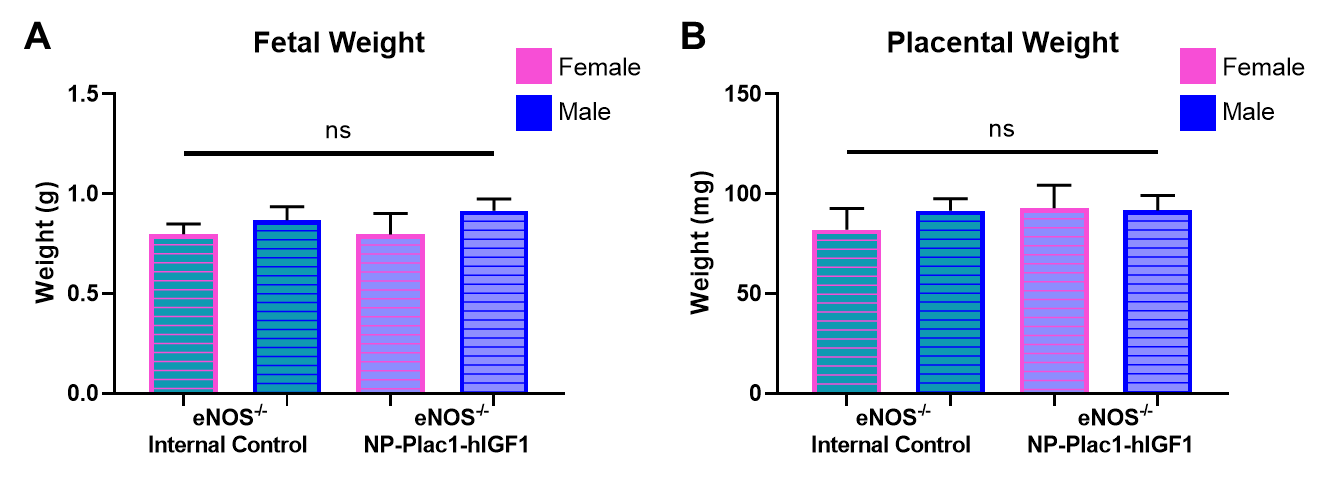

**Supplemental Figure 2** Effect of nanoparticle (NP-Plac1-hIGF1) treatment in the eNOS^-/-^ mouse placenta on fetal (**A**) and placental (**B**) weights stratified by fetal sex. There was no independent effect of fetal sex on the outcome of treatment. Data are mean±SEM. n=4 pregnant mice.

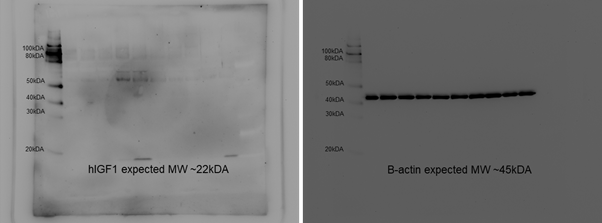

**Supplemental Figure 3** Confirmation that human insulin-like growth factor 1 (hIGF1) is not easily detected in cultured placental explants treated with nanoparticle using western blot analysis as it is a secreted protein

| **Supplemental Table S1**. Primer sequences used in qPCR analysis | | | |
| --- | --- | --- | --- |
| **Target** | **Species** | **Forward Primer** | **Reverse Primer** |
| Slc2A1 | Mouse | TGTGCTCATGACCATCGC | AAGGCCACAAAGCCAAAGAT |
| Slc2A8 | Mouse | GCCTATGTGGCAGGCTG | ATGAGCAGCAGCATGAGG |
| Slc38A1 | Mouse | GGACGGAGATAAAGGCACTC | CAGAGGGATGCTGATCAAGG |
| Slc38A2 | Mouse | TCCTTGGGCTTTCTTATGCC | TTGACACGAACGTCAAGAGA |
| Slc7A5 | Mouse | ATT CAA GAA GCC TGA GCT GG | GCA GGC CAG GAT AAA GAA CA |
| Slc7A8 | Mouse | AAGAAAGAGATCGGATTGGT | TTTCGGTGAGACAAAGATTC |
| ACTb | Mouse | GGCTGTATTCCCCTCCATCG | CCAGTTCCTAACAATGCCATGT |
| Rsp20 | Mouse | GCTGGAGAAGGTTTGTGCG | AGTGATTCTCAAAGTCTTGGTAGGC |
| IGF1 | Human | TCAGCAGTCTTCCAACCCAA | CACAGCGCCAGGTAGAAGAG |
| ACTb | Human | CGC GAG AAG ATG ACC CAG | TAG CA CAGC CTG GAT AGC AA |
| TBP | Human | GAACCACGGCACTGATTTTC | TGCCAGTCTGGACTGTTCTTC |
| BAX | Human | GGACGAACTGGACAGTAACATGG | GCAAAGTAGAAAAGGGCGACACA |
| BCL2 | Human | ATCGCCCTGTGGATGACTGAG | CAGCCAGGAGAAATCAAACAGAGG |
| Adam10 | Human | Sigma KiCqStart | |
| EPAS1 (Hif2a) | Human | Sigma KiCqStart | |
| Hif1a | Human | Sigma KiCqStart | |
| SOD1 | Human | Sigma KiCqStart | |
| SOD2 | Human | Sigma KiCqStart | |
| TP53 (p53) | Human | Sigma KiCqStart | |

| **Supplemental Table S2**. Antibodies used in immunohistochemistry | | |
| --- | --- | --- |
| **Antibody** | **Manufacturer** | **IHC Dilution** |
| Rabbit a-Slc2A1 | Abcam | 1:750 |
| Rabbit a-Slc2A8 | Abcam | 1:100 |
| Rabbit a-Slc7a5 | Abcam | 1:100 |
| Rabbit a-Slc38A2 | Abcam | 1:100 |
| Goat a-Rabbit Biotinylated Secondary | RandD | 1:200 |

| **Supplemental Table S3.** qPCR analysis of nutrient transporter mRNA in placentas from eNOS knockout (eNOS^-/-^) mice receiving nanoparticle treatment (NP-Plac1-hIGF1) | | | |
| --- | --- | --- | --- |
|  | eNOS^-/-^ Internal Control | eNOS^-/-^ NP-Plac1-hIGF1 | *P* value |
| *Slc2a1* | 1.5 (1.0-1.7) | 1.2 (1.1-1.4) | 0.5358 |
| *Slc2a8* | 1.2 (1.0-1.3) | 1.1 (1.1-1.2) | 0.8665 |
| *Slc38a1* | 1.4 (1.0-1.9) | 1.3 (1.2-1.4) | 0.6943 |
| *Slc38a2* | 1.7 (1.1-2.0) | 1.3 (1.2-1.7) | 0.6943 |
| *Slc7a5* | 1.8 (1.0-2.3) | 1.5 (1.4-1.8) | >0.9999 |
| *Slc7a8* | 2.0 (1.5-2.3) | 2.0 (1.7-2.3) | 0.6943 |
| Data are median (interquartile range) | | | |
